# INSIGHT: a population scale COVID-19 testing strategy combining point-of-care diagnosis with centralised high-throughput sequencing

**DOI:** 10.1101/2020.06.01.127019

**Authors:** Qianxin Wu, Chenqu Suo, Tom Brown, Tengyao Wang, Sarah A. Teichmann, Andrew R. Bassett

## Abstract

We present INSIGHT (Isothermal NASBA-Sequencing-based hIGH-througput Test): a two-stage COVID-19 testing strategy, using a barcoded isothermal NASBA reaction that combines point-of-care diagnosis with next generation sequencing, aiming to achieve population-scale COVID-19 testing. INSIGHT combines the advantages of near-patient with centralised testing. Stage 1 allows a quick decentralised readout for early isolation of pre-symptomatic or asymptomatic patients. The same reaction products can then be used in a highly multiplexed sequencing-based assay in Stage 2, confirming the near-patient testing results and facilitating centralised data collection. Based on experiments using commercially acquired human saliva with spiked-in viral RNA as input, the INSIGHT platform gives Stage 1 results within one to two hours, using either fluorescence detection or a lateral flow (dipstick) readout, whilst simultaneously incorporating sample-specific barcodes into the amplification product. INSIGHT Stage 2 can be performed by directly pooling and sequencing all post-amplification barcoded Stage 1 products from hundreds of thousands of samples with minimal sample preparation steps. The 95% limit of detection (LoD-95) for INSIGHT is estimated to be below 50 copies of viral RNA per 20 μl of reaction. Our two-stage testing strategy is suitable for further development into a rapid home-based and point-of-care assay, and is potentially scalable to the population level.

## Introduction

The coronavirus disease 2019 (COVID-19) pandemic is caused by the SARS-CoV-2 virus [1]. Pandemic control has been challenging due to the long incubation period and high percentage of asymptomatic carriers of this disease [2, 3]. Nucleic acid testing is thus essential to identify and isolate infected individuals at an early stage to stop the spread of the virus. At the moment, the mainstream nucleic acid test relies on a reverse transcription polymerase chain reaction (RT-PCR) assay, performed on nasopharyngeal and/or oropharyngeal swabs [4]. It requires labour-intensive RNA extraction and expensive equipment such as a thermocycler. The complexity, cost and availability of RNA extraction kits and thermocyclers have limited the throughput of such RT-PCR assays. Hence, even though the current testing regime, in conjunction with the lockdown measures, has successfully brought down the reproduction number *R*_0_ of the disease to below 1 in many countries, ramping up testing capacity sufficiently to maintain *R*_0_ below 1 will be challenging once social activities return to normal. At the same time, a prolonged lockdown is highly detrimental to the economy and the physical and mental health of individuals. Regular, high-throughput testing with rapid results is one way out of the current conundrum. Several point-of-care diagnostic tests have been proposed and some already authorised for use around the world, including SAMBA II [5], Abbott ID NOW [6] and many others. However, they typically require relatively expensive instruments or reagents, thus limiting their widespread adoption at a population level.

The ideal test would have the following five features: it would be accurate, cheap, scalable, portable and fast. This would allow for decentralised and frequent testing of a large proportion of the population, even in countries with limited medical resources. Numerous groups are working on near-patient tests [7–13] aimed at improving testing capacity and ultimately achieving regular populational scale testing. However, it is difficult to control for patient operational error and it is also challenging for centralised data collection. Centralised testing using a next generation sequencing (NGS) readout [14, 15] has also been proposed, which allows efficient scaling of testing and simple data collection, but patients do not have immediate access to the testing results thus delaying the early isolation of pre-symptomatic or asymptomatic patients. Here we propose INSIGHT (Isothermal NASBA-Sequencing-based hIGH-througput Test): a two-stage testing strategy, using a combination of isothermal Nucleic Acid Sequence-Based Amplification (NASBA) and NGS technologies, combining the advantages of near-patient and centralised testing (Figure 1a). The first stage of INSIGHT is the NASBA reaction, which can generate rapid test results on the spot in 1-2 hours. The second stage employs next generation sequencing to improve the test accuracy in a highly scalable way. Below we describe the two stages of INSIGHT in detail.

**Figure 1.**
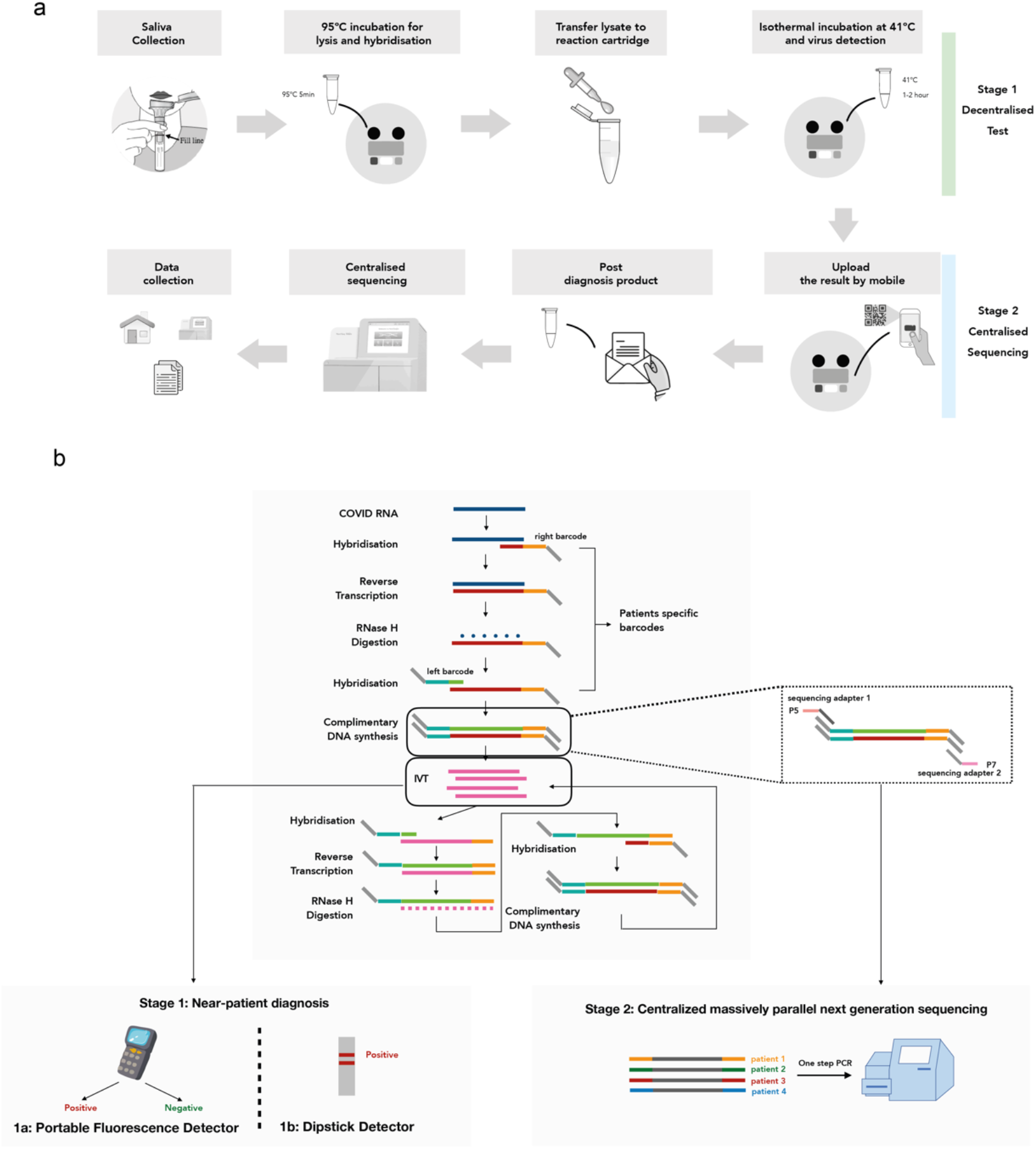
INSIGHT: an all-in-one platform for two-stage COVID-19 detection. (a) Schematic outline of our proposed two-stage testing strategy. Stage 1 is a rapid, portable decentralised COVID-19 test. Saliva can be collected directly into a tube containing QuickExtract lysis buffer at home or point-of-care. After heating up to 95 °C to lyse the virus and inactivate proteinase K, the lysate can be transferred into the NASBA reaction and incubated at 41 °C. After 1-2 hours, the result can be directly visualised with a portable fluorescence detector or lateral flow assay. In Stage 2, individuals can post the used reaction mixture to a local sequencing centre. All samples are pooled and sequenced after a one-step universal PCR. (b) Barcoded NASBA with dual readout for COVID-19 detection. In the NASBA reaction, a sample-specific left barcode is inserted between the sequencing adaptor and the forward primer, and a right barcode is inserted between the T7 promoter and the reverse primer. The NASBA *in vitro*-transcribed RNA product can be used to carry out either a portable fluorescence detection assay or a dipstick-based lateral flow assay for rapid results. Afterwards, the NASBA products can be pooled and sequenced.

The first stage consists of an isothermal NASBA reaction (Figure 1b) with crude saliva as sample input that could be incorporated into a point of care or home-based kit. NASBA employs reverse transcription and T7 RNA polymerase-mediated *in vitro* transcription to rapidly amplify RNA of interest. More than a billion-fold amplification can be typically achieved in under two hours [16]. Compared to RT-PCR, the isothermal nature of NASBA means that special equipment, such as thermocyclers, are not needed. NASBA reactions amplify both RNA and DNA, providing unique advantages for the dual stage diagnosis. For the rapid testing stage, the single stranded RNA can bind efficiently to complementary oligonucleotides without prior denaturation. A fluorescent molecular beacon can thus be used as a readout to monitor the amplification in real time and yield rapid test results [17]. Furthermore, we also established a lateral flow assay for a quick and cheap dipstick-based readout. The second stage uses NGS to further improve the test accuracy and reduce user errors in a highly scalable manner. To achieve multiplexed sequencing, a sample-specific barcode pair can be incorporated into the amplified sequence (amplicon) during the first stage reaction. The stage one end product can then be sent to a central facility for pooled sequencing, allowing up to hundreds of thousands of samples to be analysed on a single next generation sequencer. Here, the DNA in the NASBA end product is more stable and less susceptible than the RNA to degradation during the sample shipment process. NGS may also significantly reduce any possible false negative or false positive results. At the same time, the sequencing stage could be used to monitor mutations within the short amplified region, which may contribute to vaccine development and our general understanding of viral spread.

Furthermore, the INSIGHT technology can be viewed as a modular system, with the first stage consisting of two rapid test modules (either fluorescence- or dipstick-based) and the second stage a sequencing module. These modules can be used alone or combined in different ways, making INSIGHT highly flexible to adapt to different testing needs and resource availability. For example, for areas without adequate sequencing facilities, the rapid test modules (fluorescence- or lateral flow-based) in Stage 1 could be used as standalone tests. In other cases, where accessing NGS is not a limiting factor and quick assay turn-around time could be achieved by well-established logistics and NGS infrastructure, the NASBA reaction with sequencing (Stage 2) could be applied alone, reducing the need for fluorescence detectors or purchasing the lateral flow/dipstick consumables. In an ideal situation, both Stage 1 and Stage 2 could be combined, providing the benefits of both rapid and scalable diagnosis as well as centralised validation and data collection.

## Materials and reagents

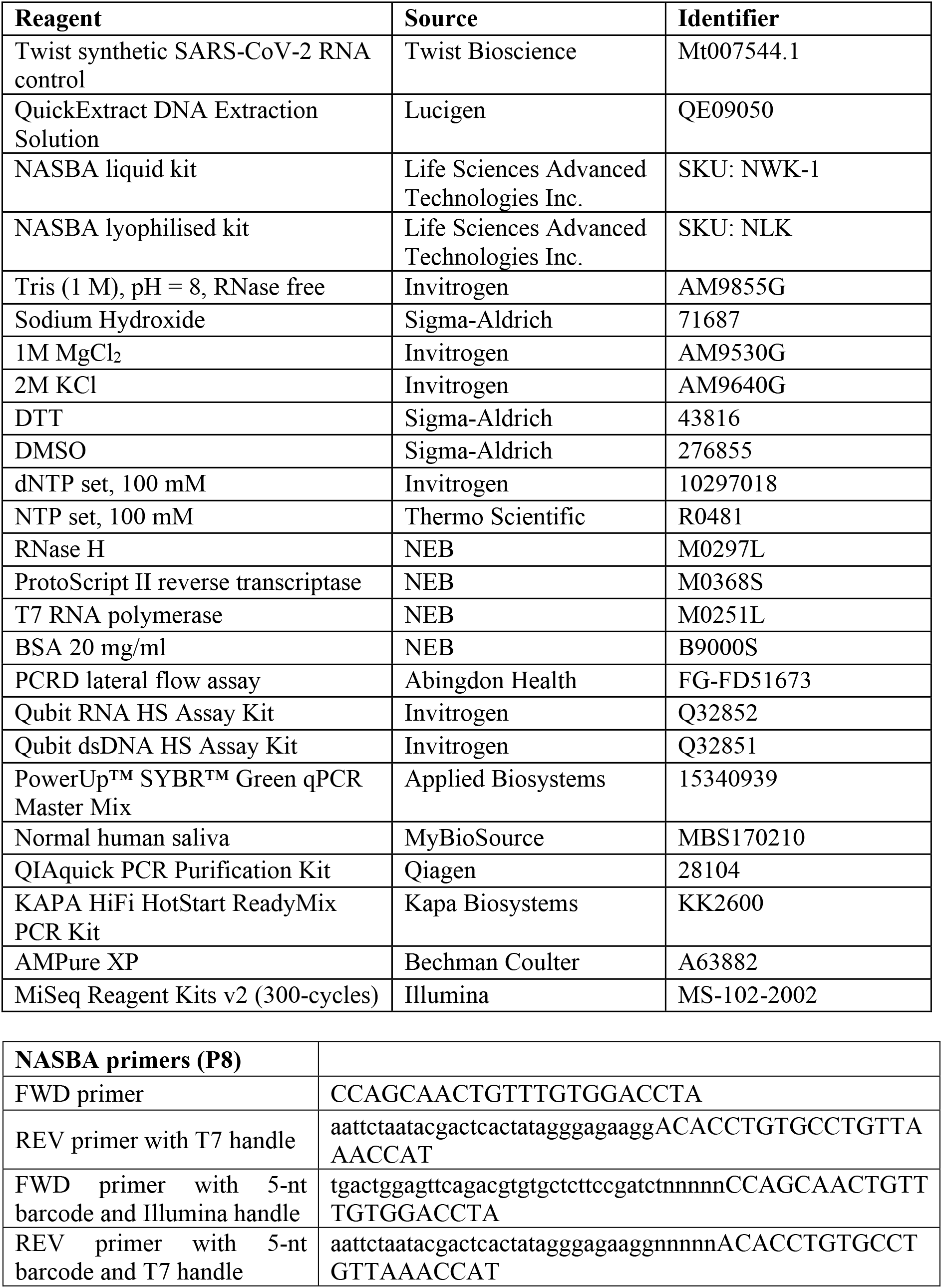
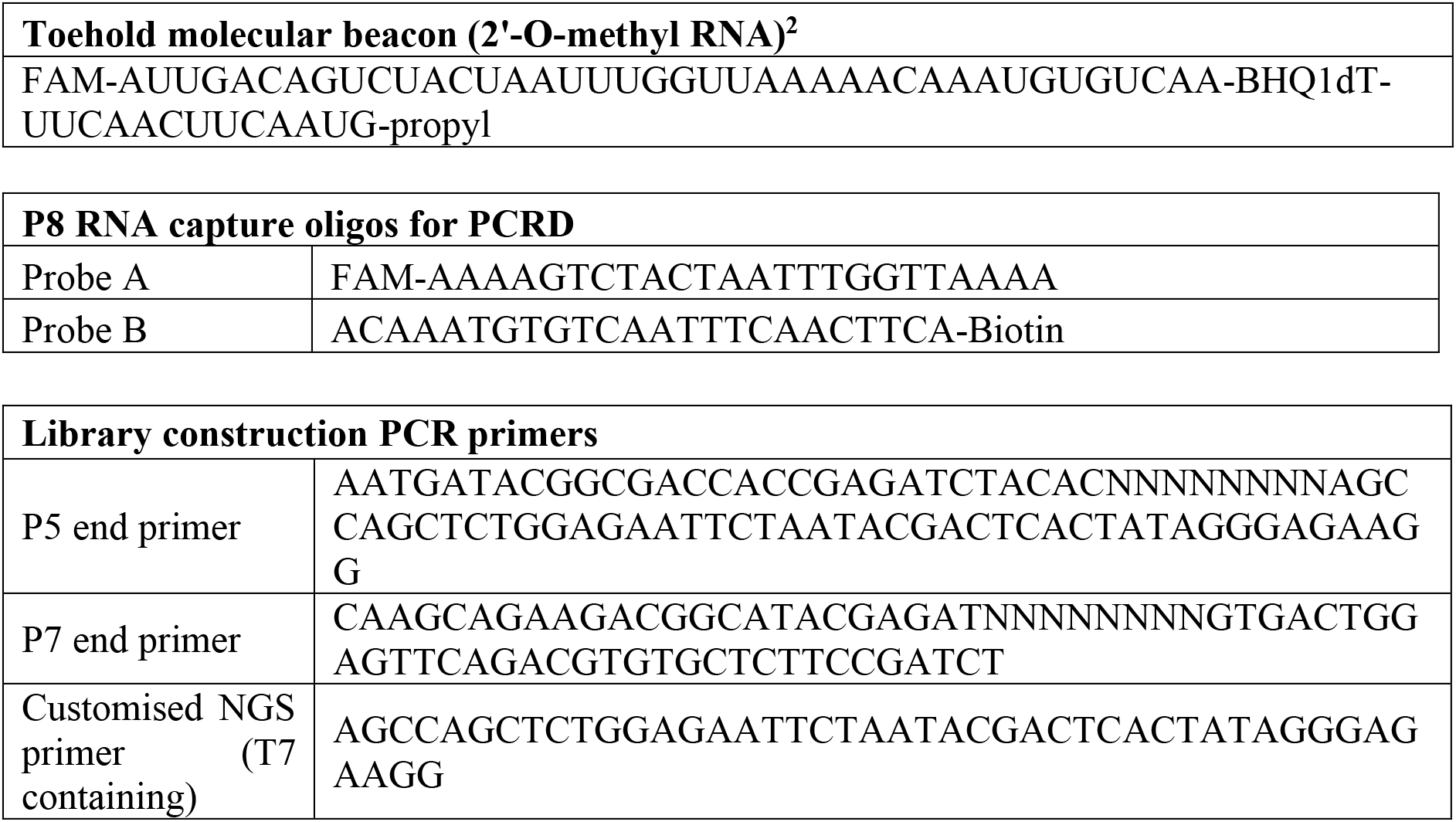

## Methods

### Step 1 Lysis of saliva samples

Mix crude saliva (commercial pooled human saliva from healthy individuals) at 1:1 ratio with QuickExtract DNA Extraction Solution. Incubate at 95 °C for 5 min to ensure complete lysis of virus and inactivation of proteinase K.

### Step 2 (Option A) NASBA reaction with fluorescence detection

Take 1 μl from the product of Step 1 (saliva lysate) and add into the NASBA reaction mixture (without the enzyme mix) to make a total volume of 15 μl. Reaction mixture can either be prepared in-house or from the Life Sciences NASBA liquid kit (see tables below) using one of the two temperature settings below.

a. Reaction mixture without the enzyme mix is incubated at 65 °C for 2 min followed by a 10-min incubation at 41 °C. Following that, 5 μl enzyme mix is added into the reaction and incubated at 41 °C for a further of 90-120 min.
b. Alternatively, reaction mixture without the enzyme mix is incubated at 95 °C for 5 min followed by a 10-min incubation at 41 °C. Following that, 5 μl enzyme mix is added into the reaction and incubated at 41 °C for a further 90-120 min.

A fluorescence plate reader (e.g. FluoSTAR) can be used to monitor the reaction in real-time, or as an endpoint assay.

*Life Sciences reaction mixture (RM)*
|  | vol. | stock conc. | conc. in RM |
| --- | --- | --- | --- |
| Saliva lysate | 1 µl |  |  |
| Primers/beacon mix | 1 µl | 500 nM each primer<br>400 nM beacon | 25 nM each primer<br>20 nM beacon<sup>3</sup> |
| Spiked-in viral RNA/water | 3 µl |  |  |
| buffer (NECB-24) | 6.7 µl |  |  |
| nucleotide (NECN-24) | 3.3 µl |  |  |
| enzyme mix (NEC-1-24) | 5 µl |  |  |
| **total volume** | 20 µl |  |  |

*In-house reaction mixture (RM)*
|  | vol. | stock conc. | conc. in RM |
| --- | --- | --- | --- |
| Saliva lysate | 1 µl |  |  |
| Primers/beacon mix | 1 µl | 500 nM each primer | 25 nM each primer |

|  |  | 400 nM beacon | 20 nM beacon |
| --- | --- | --- | --- |
| Spiked-in viral RNA/water | 4 µl |  |  |
| buffer with DMSO* | 5 µl |  |  |
| nucleotide mix* | 4 µl |  |  |
| enzyme mix* | 5 µl |  |  |
| **total volume** | 20 µl |  |  |

\*see Supplementary Table S1 for detailed composition

### Step 2 (Option B) NASBA reaction with lateral flow dipstick detection

For detection with a lateral flow assay, a NASBA lyophilised kit is used with the constitution of the reaction mixture shown below.

Take 4 μl from the product of Step 1 (saliva lysate) and add into the NASBA reaction mixture (without the enzyme mix) to make a total volume of 60 μl. Incubate at 95 °C for 5 min followed by a 10-min incubation at 41 °C.

Following that, 20 μl enzyme mix is added into the reaction and incubated at 41 °C for a further of 90-120 min. Take the reaction product to the sample well of a PCRD test cassette. Results will be shown within 10 min.

|  | **vol.** | **stock conc.** | **conc. in RM** |
| --- | --- | --- | --- |
| Saliva lysate | 4 µl |  |  |
| primers | 4 µl | 500 nM each primer | 25 nM each primer |
| Spiked-in viral RNA/water | 8.16 µl |  |  |
| Probe A | 1.92 µl | 1 µM | 24 nM |
| Probe B | 1.92 µl | 1 µM | 24 nM |
| Reconstituted buffer mix (LRB) | 40 µl |  |  |
| Reconstituted enzyme mix (LEM) | 20 µl |  |  |
| **total volume** | 80 µl |  |  |

### Step 3 Library construction for NGS

To allow for pooled sequencing of NASBA reaction end products, barcode sequences are added upstream of each of the forward and reverse primers (Figure 3a). In addition, an Illumina sequencing adaptor is added upstream of the forward primer barcode sequence as a universal PCR handle (see Materials and reagents section for the exact sequence).

Here, 2 μl NASBA end products from each sample are first pooled into a single tube. Pooled products are then column purified to remove residual NASBA primers (QIAquick PCR Purification Kit). PCR is performed on the column purified pooled sample using two NGS indexing primers. Here, we have designed a customised NGS primer containing the T7 polymerase promoter sequence (see Materials and reagents section for the exact sequence) at the P5 end and used a standard TruSeq sequencing primer at the P7 side. A PCR mix is made based on the table below. A standard PCR program is used with longer elongation time and minimal cycle number to reduce barcode hopping.

|  | **vol.** |
| --- | --- |
| 2x PCR mix (KAPA HIFI HotStart ReadyMix) | 20 µl |
| P5 end primer (10 µM) | 1 µl |
| P7 end primer (10 µM) | 1 µl |
| Column purified NASBA product (~3.5 ng dsDNA) | 4 µl |
| Nuclease free water | 14 µl |

| **temperature** | **time** | **cycle number** |
| --- | --- | --- |
| 95 °C | 3 min | 1 |
| 98 °C | 20 s | 15 |
| 60 °C | 15 s |  |
| 72 °C | 30 s |  |
| 72 °C | 4 min | 1 |

After the PCR, an AMPure bead-based double size selection is carried out (0.55x and 0.75x) to enrich for products of interest. In this study, a MiSeq Reagent Kit v2 (300-cycles) was used for NGS.

### Step 4 Analysis of NGS results

To analyse the INSIGHT NGS data, sequences in FASTQ files are first trimmed to leave the first 80 nucleotides for both read 1 and read 2 using FASTX_trimmer. The trimmed read 1 and paired read 2 are then merged by FLASH. The merged sequence is compared with the reference viral genome sequence (NNNNNACACCTGTGCCTGTTAAACCATTGAAGTTGAAATTGACACATTTGTTTTTAACCAAATTAGTAGACTTTTTAGGTCCACAAACAGTTGCTGGNNNNN, N stands for the barcode position), and only those with a hamming distance less than or equal to 2 are extracted. Here, only substitutions were allowed while insertion- and deletion-containing reads were filtered out. The first 5 nt and the final 5 nt regions of all extracted sequences correspond respectively to the right barcode and the reverse complement of the left barcode. Diagnostic results for sequenced NASBA samples are determined according to the read counts of their corresponding sample-specific barcode pairs. More details can be found in the Results section.

## Results

### INSIGHT technology development and optimisation

Primers were designed to target the SARS-CoV-2 S gene, which encodes the viral envelope spike glycoprotein. The S gene is one of the most highly expressed viral RNAs [18], and at the same time is a promising target for SARS-CoV-2 vaccine development [19]. In addition, the S gene sequence is an informative sentinel for viral genome evolution with respect to increased or decreased affinity to the human viral entry receptors such as ACE2 [20]. We screened 13 pairs of primers and selected the most efficient pair to optimise our in-house NASBA reaction (Supplementary Figure S1a). All subsequent reactions and figures shown in this paper were carried out with primer pair P8 (sequence shown in Materials and reagents). With synthetic SARS-CoV-2 RNA as the input, we have achieved target RNA amplification with both our in-house NASBA mixture and a commercial ready-made mixture. The NASBA reaction product was confirmed by an RNA urea gel (Supplementary Figure S1b), showing a product of the expected size in the presence of viral template RNA only. Furthermore, we also optimised the primer concentration used in the NASBA reaction by quantifying the NASBA end product RNA concentration with a molecular beacon (Supplementary Figure S1c). We found a primer concentration of 25 nM each, which is 10 times less than typically used in the literature [21–23], increased the reaction efficiency. Additional optimisation has been carried out for our in-house NASBA reaction mixture. This includes the choice of enzymes (ProtoScript II reverse transcriptase instead of AMV reverse transcriptase, Supplementary Figure S1d), the concentration of enzymes (higher concentration with better yield and more consistent results, Supplementary Figure S1e) and the buffer pH (Supplementary Figure S1f).

To work towards an INSIGHT test that can be used at home, and to simplify the assay workflow and improve its scalability, the NASBA reaction would ideally be applied to a crude saliva lysate. This would alleviate the need for complex and expensive processes to purify RNA. Commercially acquired human saliva can be mixed with QuickExtract buffer and heat treated at 95 °C for 5 min to generate saliva lysate [24]. Here, we demonstrate that saliva lysate is compatible with the NASBA reaction using commercially available human saliva from healthy individuals with spiked-in synthetic viral RNA. To minimise the handling steps, saliva lysate is combined with partial NASBA mix (without the enzyme cocktail) and heated at 95°C for 5 min to inactivate the proteinase K and disrupt RNA secondary structure (Supplementary Figure S2a). This suggests that in practice, the 95 °C denaturation step could be combined with the 95 °C viral lysis step to save an extra heating step. The method performs well, with 100 copies of input RNA reliably detected.

Finally, to achieve better assay sensitivity, we also varied saliva lysate input volume to find the maximum compatible saliva input amount. We found saliva lysate can be increased from 1 μl to 3 μl in a total reaction volume of 20 μl, and the detection threshold was not compromised (Supplementary Figure S2b).

### INSIGHT Stage 1 (Option A): fluorescence readout with molecular beacon detection

We have established two forms of specific readouts for Stage 1 of INSIGHT: a molecular beacon with fluorescence detection, and a dipstick-based lateral flow assay. We describe the performance of the former option here and the latter in the next subsection.

Molecular beacons are hairpin-shaped molecules with a fluorophore and a quencher covalently attached and brought into close proximity due to the secondary structure formed by the hairpin. Upon recognition of the target, the fluorophore and the quencher will be spatially separated due to hybridisation, which results in fluorescence (Figure 2a). Here, four types of beacons were designed to target the P8 amplicon: a conventional DNA beacon, a toehold DNA beacon, a conventional 2’-O-methyl RNA beacon and a toehold 2’-O-methyl RNA beacon. The toehold provides an initial anchor point for the beacon to latch onto its target and assists in the unwinding of the stem of the beacon, and the 2’-O-methyl modification increases target affinity and provides stability against oligonucleotide degradation. All four types were first tested with *in vitro* transcribed RNA, and the 2’-O-methyl RNA toehold beacon was found to achieve the best sensitivity (Figure 2b). Therefore, we chose the RNA toehold beacon for our COVID-19 NASBA assay. An essential feature of this beacon is a 3’-propyl group which prevents any possibility of polymerase extension of the toehold sequence.

**Figure 2.**
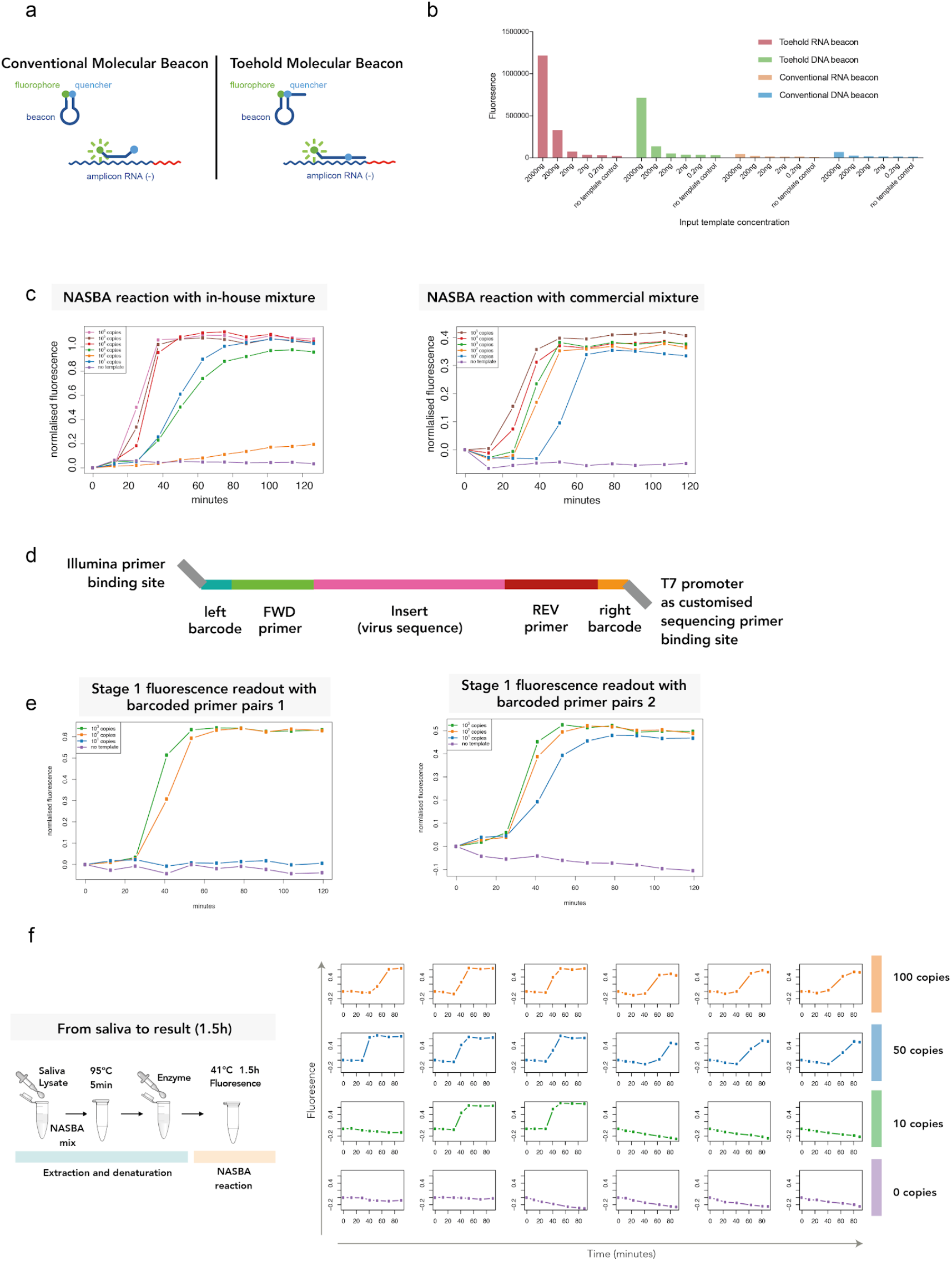
INSIGHT Stage 1 fluorescence readout with molecular beacon detection. (a) Illustration of conventional and toehold molecular beacons. (b) Sensitivity of different types of molecular beacons. Four different types of beacons were mixed with increasing amounts (0.2 ng to 2000 ng) of *in vitro*-transcribed target amplicon RNA for a fluorescence readout. (c) In-house and commercial NASBA mixtures with varying amounts (10^1^ to 10^6^ copies) of input viral RNA templates and a real-time molecular beacon readout. With a final concentration of 20 nM of beacon in the reaction, fluorescence was detectable at around 30 mins, and reached saturation at around 60 mins. (d) Schematic illustration of the barcoded NASBA DNA product. Using the primer pair P8, the forward primer contains a 24-nt sequence targeting part of the SARS-CoV-2 S gene, a 5-nt left barcode and a 33-nt sequence of the Illumina sequencing adaptor. The reverse primer contains a 22-nt sequence complementary to part to SARS-CoV-2 S gene, a 5-nt right barcode and a 31-nt T7 promoter. Customised sequencing primers can be used for the reverse read. (e) With barcoded primers, the NASBA reaction achieved a detection threshold of around 10-100 copies of viral RNA per 20 μl reaction for the commercial NASBA mixture (2 replicates). (f) Left: Schematic illustration of experimental setup with 5 min at 95 °C to denature the viral RNA, followed by 1.5 hr at 41 °C for the NASBA reaction. Right: Real-time INSIGHT Stage 1 fluorescence readings for samples with 0, 10, 50 and 100 copies of viral RNA per 20 μl reaction (6 technical replicates each).

We have also titrated the concentration of the toehold RNA beacon in the reaction mixture to achieve the best real-time results, as a high beacon concentration inhibits the NASBA reaction (Supplementary Figure S1g). We found that 20 nM of molecular beacon resulted in the best assay sensitivity. With this beacon concentration, the fluorescence plateaus at around 60 min into the reaction. With the non-barcoded primer pair P8, the real-time detection limit is around 100-1000 copies/reaction for our in-house mixture, and around 10 copies/reaction for the commercial mixture (Figure 2c).

In order to incorporate patient specific barcodes in the NASBA reaction, we have designed the primers with 5-nt barcodes flanked on each side, and an Illumina handle on the forward primer (sequences shown in Materials and reagents) so that the NASBA DNA product would contain a 5-nt barcode at each end (Figure 2d). The NASBA reaction still worked with the barcoded primers, despite a slightly reduced detection threshold of around 10-100 copies/reaction for the commercial NASBA mixture (Figure 2e). By using combinatorial barcoding, a 5-nt barcode sequence on both sides can generate up to a million unique barcodes, or 30,000 Hamming-distance 3-separated barcodes. The latter allows single nucleotide substitution error in the barcodes to be corrected after sequencing. If needed, a 6-nt barcode on each primer can generate 450,000 Hamming-distance 3-separated unique barcode pairs. (The software code for barcode generation is available at: *www.github.com/suochenqu/INSIGHT*)

We assessed the performance of INSIGHT Stage 1 with real-time fluorescence detection using human saliva from healthy individuals with spiked-in synthetic viral RNA, following temperature setting (b) in Step 2A of the Methods section (Figure 2f). For samples with input of 50 viral RNA copies per 20 μl reaction, all 6 reactions showed amplification of fluorescence signals, whereas for samples with input of 10 viral RNA copies per 20 μl reaction, 2 out of 6 reactions showed amplification.

### INSIGHT Stage 1 (Option B): dipstick readout with lateral flow assay

We have also developed and optimised a dipstick-based lateral flow assay as an alternative to fluorescence detection in Stage 1 of INSIGHT. Compared to molecular beacon detection where a portable fluorescence detector is needed, dipstick-based detection is a particularly attractive option as it does not require any extra equipment apart from a portable heating source.

Here, we use a format of lateral flow assay which detects nucleic acid utilising neutravidin-conjugated carbon nanoparticles (NA-CNPs) that can bind to biotin, a test line comprising an anti-FAM antibody and a biotin control line to capture excess NA-CNPs. Two RNA capture oligos are added into the NASBA reaction. One is FAM-labelled and the other is biotin-labelled. During the reaction, both capture oligos bind to different parts of the single-stranded RNA NASBA product. The dual tagging of FAM and biotin results in aggregation of NA-CNP at the test line, resulting in a visible signal in the assay (Figure 3a). A positive lateral flow assay would show a clearly visible line at test line 2 in addition to the control line (C-line) whereas only the C-line is visible in a negative assay.

**Figure 3.**
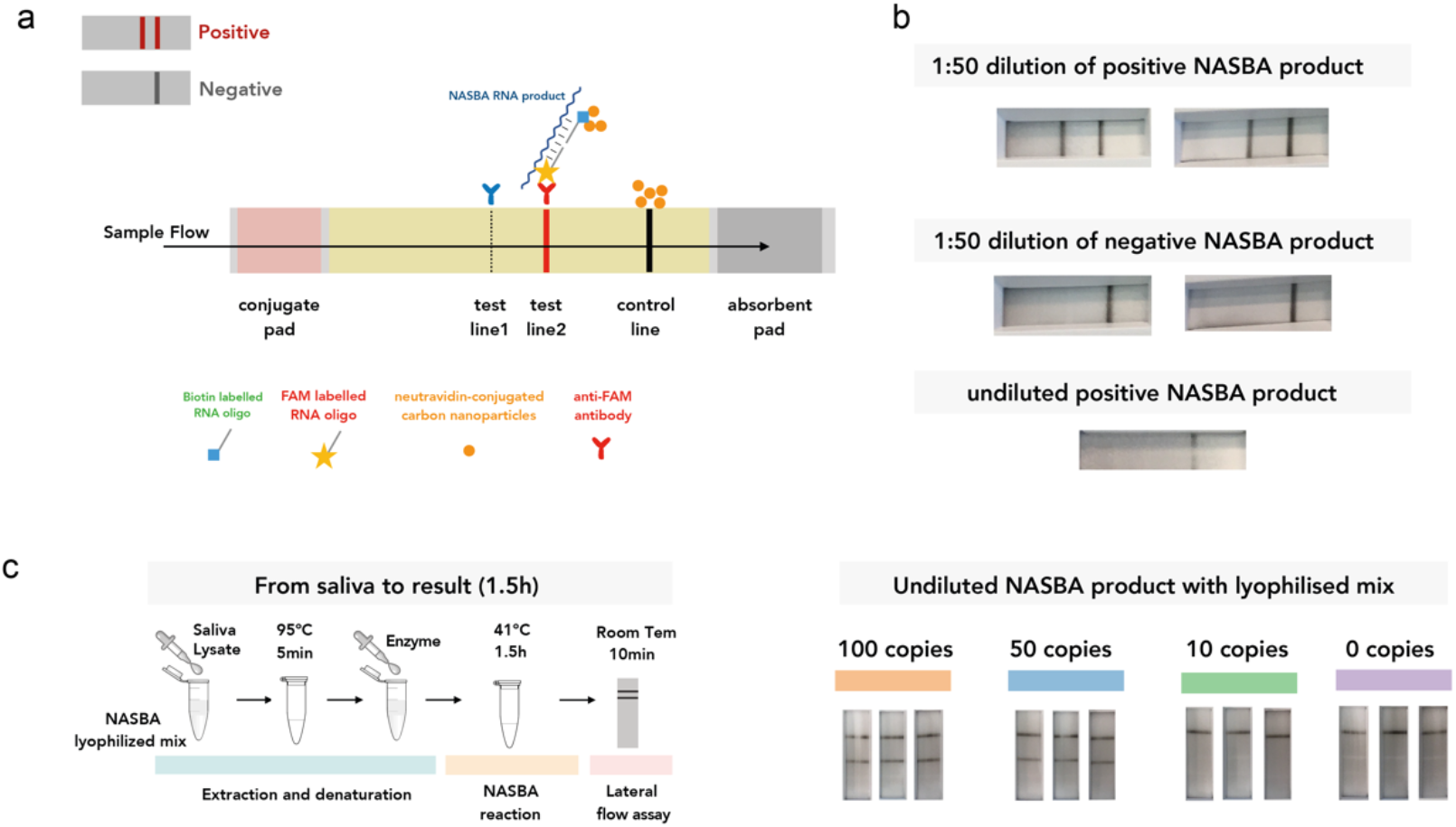
INSIGHT Stage 1 dipstick readout with lateral flow assay. (a) Schematic illustration of the nucleic acid lateral flow assay. (b) Lateral flow-based dipstick test results from NASBA reactions using liquid NASBA mix. The lateral flow assay works when the NASBA end product is diluted 50 times with PCRD buffer, but not with undiluted NASBA end product. (c) Left: Schematic illustration of experimental setup with 5 min at 95 °C to denature the viral RNA, followed by 1.5 hr at 41 °C for the NASBA reaction. Lyophilised NASBA mix is used in the reaction. The NASBA end product can be directly loaded onto the dipstick. Results can be read after 10 minutes incubation at room temperature. Right: All three technical replicates for 50 copies viral RNA per 20 μl reaction were detectable using the dipstick assay.

By using the commercially available lyophilised NASBA mix, samples at the end of the NASBA reaction can be directly loaded onto the dipstick without any extra step of RNA purification or dilution (see Methods section Step 2 Option B for details).

We also assessed the performance of INSIGHT Stage 1 with dipstick detection using human saliva from healthy individuals with spiked-in synthetic viral RNA (Figure 3c). Three technical repeats were performed for different amounts of viral RNA input. This successfully detected viral RNA with input of 50 copies per 20 μl reaction, and failed to detect samples with input of 10 copies per 20 μl reaction. We used barcoded primers in the reactions here to make the Stage 1 product compatible with Stage 2 sequencing.

We note that it is necessary to use the lyophilised mix to achieve a sample-to-dipstick result without any extra steps. We have also tried using the commercially available liquid NASBA mix, but found that it required an extra step of dilution prior to the lateral flow assay (Figure 3b), possibly due to a component within the liquid mix interfering with the downstream lateral flow assay. In addition, we previously attempted to use biotinylated UTP and FAM-tagged RNA capture oligo for dual tagging. However, this combination requires an extra step of RNA purification after the NASBA reaction to prevent unused biotinylated UTP from saturating the NA-CNPs (Supplementary Figure S3). In summary, we have optimised the conditions such that the lyophilised mix allows direct sample-to-result readout.

### INSIGHT Stage 2: pooled library construction and NGS readout

After the NASBA reaction, which can occur in a near-patient setting, we propose that the INSIGHT Stage 1 products can be transported to a local sequencing centre. As sample-specific barcodes would have already been incorporated in Stage 1 in a decentralised manner, all samples can be directly pooled, and the NGS library prepared, with a simple one-step PCR using primers flanked with sequencing adapters (Figure 4a, see Methods section Step 3 for details).

**Figure 4.**
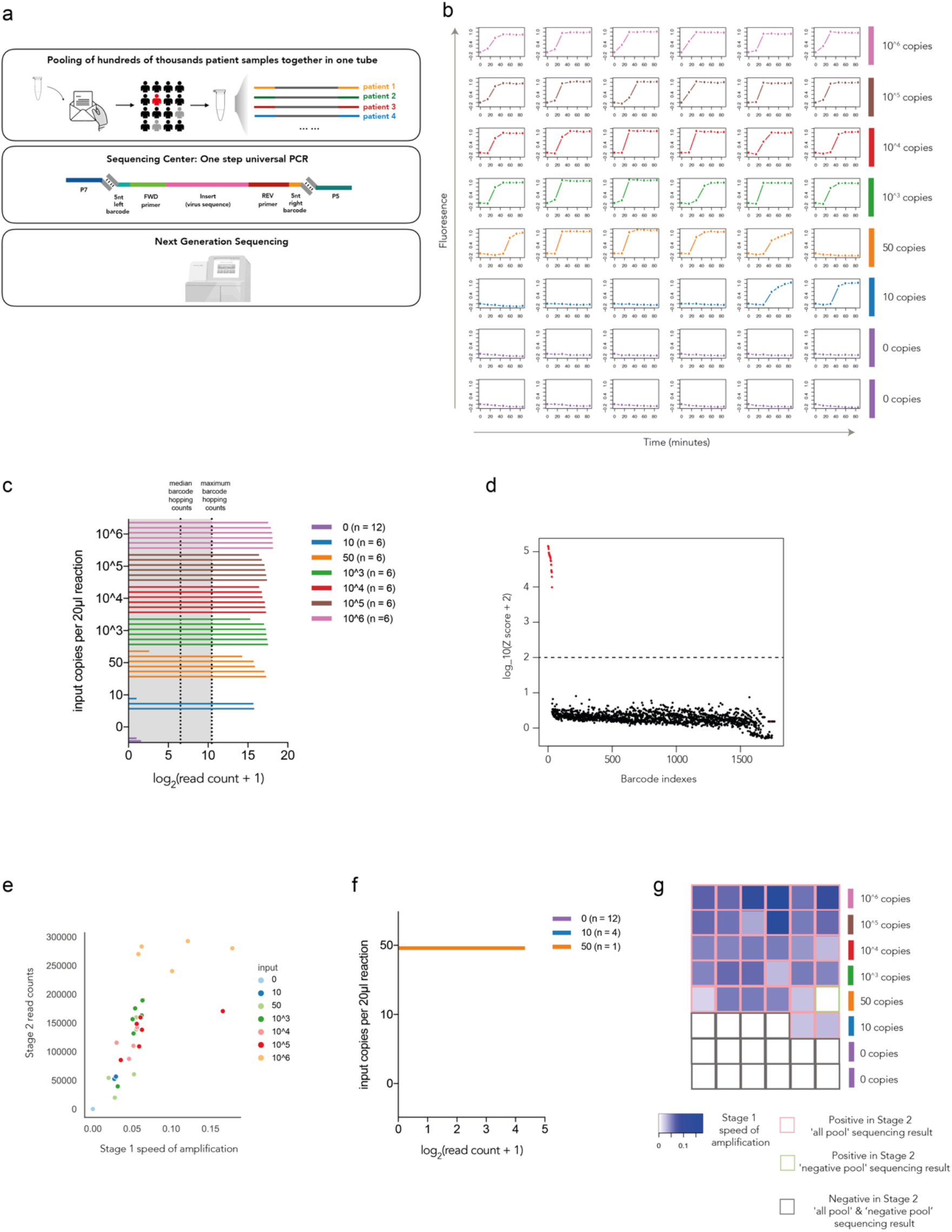
INSIGHT Stage 2 pooled testing using next generation sequencing. (a) Strategy for Stage 2 NGS-based detection. Up to hundreds of thousands of samples can be pooled together, and a universal indexing PCR can be carried out at a local sequencing centre. (b) Real-time molecular beacon readings for 48 barcoded primer pairs with various amounts (0 to 10^6^ copies) of viral RNA input per 20 μl reaction. (c) INSIGHT Stage 2 read counts for samples from all reactions in (b) i.e. ‘all pool’. The two dotted lines show the median and the maximum of barcode hopping read counts. (d) The Z-score distribution of used barcode pairs and hopped barcode pairs using statistical modelling. Red dots: used barcode pairs. Black dots: hopped barcode pairs. (e) Correlation analysis of INSIGHT Stage 1 speed of amplification (reciprocal of time to reach 0.2 normalised fluorescence reading) and Stage 2 sequencing read counts. (f) INSIGHT Stage 2 read counts for samples from 17 reactions in (b) that showed negative results in Stage 1 fluorescence detection i.e. ‘negative pool’. (g) Combined results from both Stage 1 and 2. The colour of the heatmap represents INSIGHT Stage 1 speed of amplification with the same sample order in (b). Samples that were positive in INSIGHT Stage 2 ‘all pool’ sequencing results are boxed in red. Samples that were positive in INSIGHT Stage 2 ‘negative pool’ sequencing results are boxed in green. Samples that were negative in both ‘all pool’ and ‘negative pool’ sequencing results are boxed in black.

We designed an experiment with 48 contrived samples (Figure 4b). Each reaction used a unique pair of barcoded primers. We varied the input of the viral RNA as shown in Figure 4b to mimic the wide range of viral load in patient samples. The first stage of INSIGHT was carried out with real-time fluorescence detection. For samples with input of 50 viral RNA copies per 20 μl reaction, 5 out of 6 reactions showed amplification of fluorescence signals, whereas for samples with input of 10 viral RNA copies per 20 μl reaction, 2 out of 6 reactions showed amplification (Figure 4b). For the second-stage sequencing, we performed two pooling strategies either independent of the first-stage results or dependent on the first-stage results. ‘All pool’ included all 48 samples regardless of the first-stage results. ‘Negative pool’ was the collection of the 17 samples that showed negative results in the first stage. Library preparation was performed separately for ‘all pool’ and ‘negative pool’.

Figure 4c shows the read counts from all samples in ‘all pool’. In addition to reads that contain the expected barcode pairs, we also observed some reads with left and right barcode combinations that were not used in the experiment, a phenomenon we refer to as ‘barcode hopping’. The two dotted lines in the plot of Figure 4c indicate the median and the maximum read counts of all hopped barcode pairs. Read counts from samples with 50 viral RNA copies and 10 viral RNA copies showed that 5 out of 6 and 2 out 6 respectively had higher read counts than maximum read count of hopped barcode pairs, providing an identical result to the fluorescent readout. In this particular experiment, both left and right barcodes in every sample are unique, *i.e.* no two samples share the same left or right barcode. Therefore, hopped barcode pairs can be unambiguously identified and evaluated in this experiment.

In order to increase the available barcode pair combinations for multiplexing, repeated usage of the same left or right barcode is desirable. We have thus sought to evaluate and potentially circumvent the barcode hopping problem, and have built a statistical model (Supplementary Note) to predict the number of reads generated from barcode hopping. If the observed read count of a barcode pair is significantly larger than the predicted read count (Z score exceeding 100), then the barcode pair is likely to be a real product of NASBA amplification, *i.e.* a positive result. Using the above calling procedure, we can unambiguously identify 31 out of the 36 samples with viral RNA input as ‘positive’ and all 12 negative control samples as ‘negative’ in ‘all pool’ (Figure 4d). This result is identical to the Stage 1 fluorescence result.

We also performed a correlation analysis between the INSIGHT Stage 1 speed of signal amplification (measured as the reciprocal of time to reach a normalised fluorescence level of 0.2) and the Stage 2 read counts. As expected, there is a strong positive correlation between the results from the two stages (Pearson R = 0.67, p < 0.0001, see Figure 4e). Interestingly, although the input RNA varied from 10 to 10^6^ copies per reaction, the INSIGHT Stage 2 read counts for amplified samples only differed by 14-fold. This showed that the NASBA reaction in Stage 1 had saturated for most samples, thus allowing Stage 2 to handle samples across an extremely wide dynamic range of viral RNA input molecules.

Figure 4f shows the read counts from all samples when sequenced in the ‘negative pool’. One sample, which had 50 copies of viral RNA as input and failed to be picked up as ‘positive’ in Stage 1, showed positive read counts in Stage 2. This means that Stage 2 sequencing can further improve the sensitivity of Stage 1 results. INSIGHT Stage 1 and Stage 2 combined results are summarised and shown in Figure 4g.

### INSIGHT performance evaluation and comparison with RT-qPCR

We next sought to summarise the limit of detection (LoD) for the INSIGHT technology. Here LoD-95 was defined to be the input viral RNA amount at which 95% of samples can be detected. This is estimated by maximum likelihood estimation under the assumption that the number of viral RNA copies in the reaction input follows a Poisson distribution, and that each molecule has the same probability of being amplified. INSIGHT Stage 1 with a fluorescence readout attains an estimated LoD-95 of 63.7 (CI = [36.8, 113]) copies per 20 μl reaction (Figure 5b). This is calculated based on the 24 reactions shown in Figure 2f and the 48 reactions shown in Figure 4b. Separately, INSIGHT Stage 1 with dipstick readout has an LoD-95 of 75.8 (CI = [24.9, 234]) copies per 20 μl reaction, which is computed using reactions shown in Figure 3c. For the second stage, using the ‘all pool’ results shown in Figure 4c, we calculated that the NGS sequencing alone has a LoD-95 of 80.3 copies (CI = [37.7, 197]) per 20 μl reaction. By combining the Stage 1 fluorescence readout and Stage 2 ‘negative pool’ results, the overall INSIGHT technology LoD-95 can be further improved to 49.9 (CI = [21.2, 125]).

**Figure 5.**
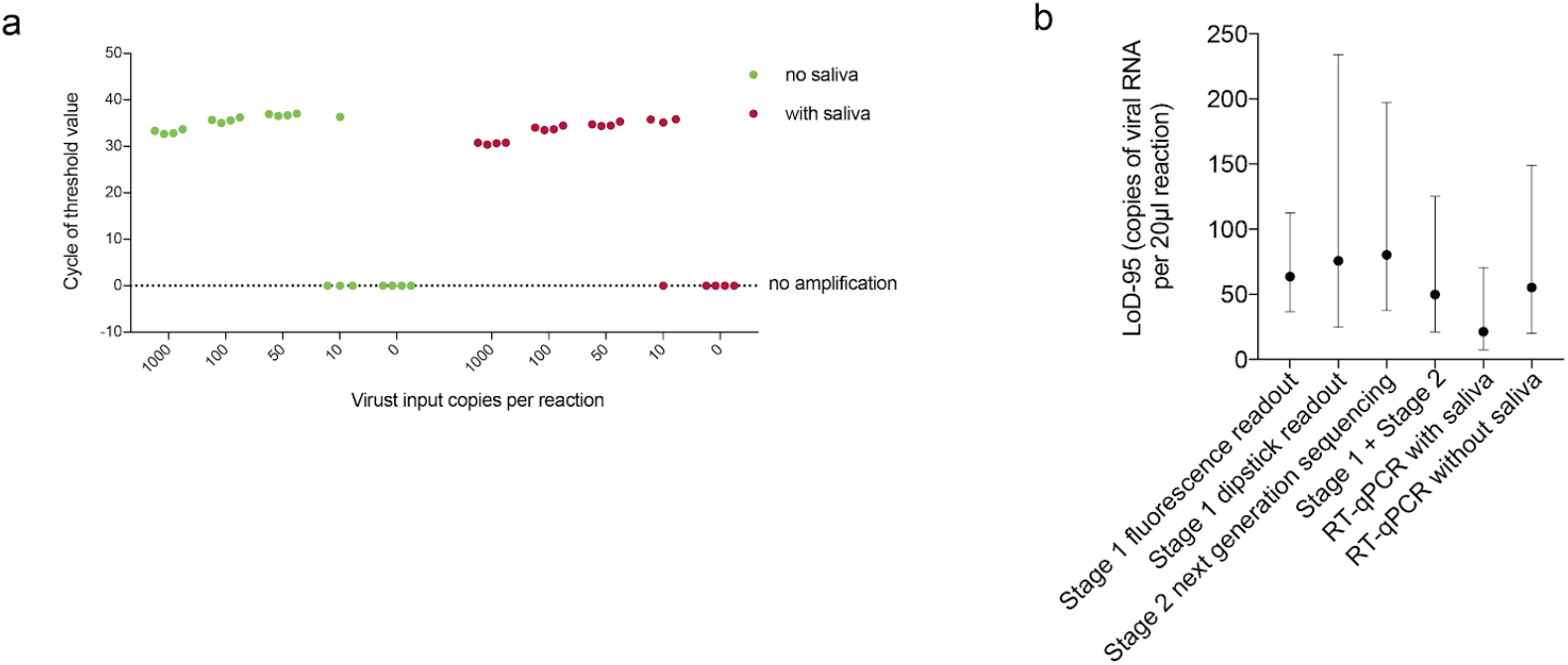
INSIGHT technology limit of detection and comparison with RT-qPCR. (a) The qPCR cycle number to reach fluorescence threshold (Ct) using the Public Health England protocol and various numbers of viral RNA copies per 20 μl reaction as input. (b) The estimated 95% limit of detection (LoD-95) for different stages of INSIGHT and gold standard RT-qPCR, with the whiskers spanning the 95% confidence intervals of the estimates.

To put our LoD figures in context, we compared our method with the gold standard RT-qPCR assay. We performed RT-qPCR (Figure 5a) following the protocol recommended by the Public Health England (PHE) in England [25]. Chemically synthesised viral RNA was used as sample input. Four technical replicates each were carried out with and without saliva lysate from healthy individuals. The RT-qPCR could consistently detect samples with 50 copies of viral RNA per 20 μl reaction. With 10 copies of viral RNA per 20 μl reaction, it detected 3 out of 4 samples with saliva lysate and only 1 out of 4 without saliva lysate. Thus, the estimated LoD-95 for the RT-qPCR is 55.3 (95% confidence interval = [20.2, 149]) copies per 20 μl reaction without saliva, and 21.4 (CI = [7.3, 70.5]) copies per reaction with 1 μl saliva lysate added in the reaction. In summary, the INSIGHT protocol is highly comparable to the PHE RT-qPCR protocol in terms of sensitivity.

## Discussion

INSIGHT’s dual-stage design combines the benefits of both near-patient testing and centralised testing. It offers a rapid first stage readout without delay and minimises the problem of RNA degradation. The distributed first stage also reduces the logistic burden and labour requirements for carrying out population scale screening. The centralised second stage can be used to eliminate user errors, collate results in a centralised repository and help inform other epidemiological efforts. In addition, INSIGHT’s two stages can be viewed as three different modules – two rapid detection modules (fluorescence detection or dipstick-based detection), and one sequencing module. The modules can be either combined in a two-stage test as illustrated here, or used independently. This offers flexibility for adaptation to local needs and resources, and for testing other viruses.

Our assay sensitivity is comparable to the gold standard RT-qPCR test. Although INSIGHT has shown promising results in experimental settings, additional work is required to bring it into practical use. Most importantly, the current experiments were carried out using saliva from healthy individuals with spiked-in viral RNA rather than patient samples.

In addition, several new techniques we developed as part of the INSIGHT technology may be of interest in other contexts. First, we show that toehold beacons can achieve a much higher sensitivity than commonly used regular molecular beacons for fluorescence detection. Secondly, the use of two RNA capture oligos with different tags in the INSIGHT Stage 1 dipstick readout offers a new way to detect RNA in a lateral flow assay, and further increases specificity for a particular target sequence. Finally, we have established a new statistical model to address the problem of barcode hopping that can be used in any multiplexed NGS settings.

Other isothermal methods have been proposed for SARS-CoV-2 detection. Most of them are based on the RT-LAMP technology with a colourimetric or turbidimetric readout [7–10]. These assays could potentially suffer from false positive results generated from nonspecific primer binding or primer dimers [11]. To circumvent this, [12, 13] have proposed to use a CRISPR-based assay to achieve highly specific test results. A major advantage of INSIGHT is to enable a rapid readout with high specificity for SARS-CoV-2, whilst avoiding complex additional steps such as CRISPR-based cleavage. In the first stage, a molecular beacon for fluorescence readout or RNA capture oligos for lateral flow readout provide an additional layer of sequence-specific detection. In the second stage, NGS further improves the assay sensitivity and specificity by unambiguously identifying the viral sequence. We note that, unlike the concatemer amplification product in LAMP, the NASBA reaction generates a single, well-defined amplicon, thus making it particularly suitable for sequencing. Furthermore, it allows combinatorial barcoding, thus avoiding the need to synthesise hundreds of thousands of barcoded oligonucleotides.

While different research groups [14, 15] and companies (e.g. COVIDSeq [26] from Illumina and SwabSeq [27] from Octant) have proposed to use NGS to expand testing capacity, most of them are RT-PCR-based, restricting amplification and barcoding to centralised facilities. The decentralised first stage NASBA reaction in INSIGHT greatly reduces the burden of sample handling in testing centres, and hence makes regular population-scale screening feasible. Other groups including [11, 28–30] have also independently suggested similar approaches of barcoded isothermal amplification.

Our system can also be modified to include additional primers targeting a different region of the SARS-CoV-2 genome or other pathogens. In future, a positive control RNA sequence that can be amplified by one of the same primer pairs can be added to the reaction, to ensure a negative result is not due to faulty reagents or patient handling mistakes, and provide some degree of quantification of viral load.

The global COVID-19 emergency has exposed our weaknesses in responding to a new pathogen and an emerging pandemic, and we hope the INSIGHT two-stage testing strategy has potential to impact this pandemic and beyond.

## Acknowledgement

We thank D. Kwiatkowski, C. Langford, M. Lawniczak, M. Dougherty and the CellGen admin team (all Wellcome Sanger Institute) for supporting this project. We also thank M. Chhatriwala, S. Gozna, M. Hurles, A. Ibrahim, S. Palmer and C. Smee (all Wellcome Sanger Institute) for their helpful comments on an earlier version of this manuscript. We are grateful to A. Peltan, H. Chick, F. Euan from New England BioLabs, G. Grassmann, C. Mihm, F. Perez from Life Sciences Advanced Technologies Inc., A. Lightowlers, D. JHF Knapp from Oxford University for helpful discussions. We acknowledge funding from the Wellcome Project Heron and the Wellcome Sanger Institute core grant from Wellcome [WT206194]. C.S. is supported by a Wellcome Trust PhD Fellowship for Clinicians.

## Declaration of conflicts of interest

Q.W., C.S., S.A.T. and A.R.B. are inventors on a patent application (GB2007724.4) submitted by Genome Research Limited, with the aim of making this technology freely available for research and deployment. In the past three years, S.A.T. has consulted for Genentech and Roche, and sits on Scientific Advisory Boards for Foresite Labs, Biogen and GlaxoSmithKline. T.B. is a consultant at ATDBio.

## Data and material availability

This protocol is also available at: www.protocols.io/view/insight-a-scalable-isothermal-nasba-based-platform-bghsjt6e.

The software code for barcode generation and barcode hopping analysis is available at: www.github.com/suochenqu/INSIGHT.

## Supplementary Information

**Figure S1.**
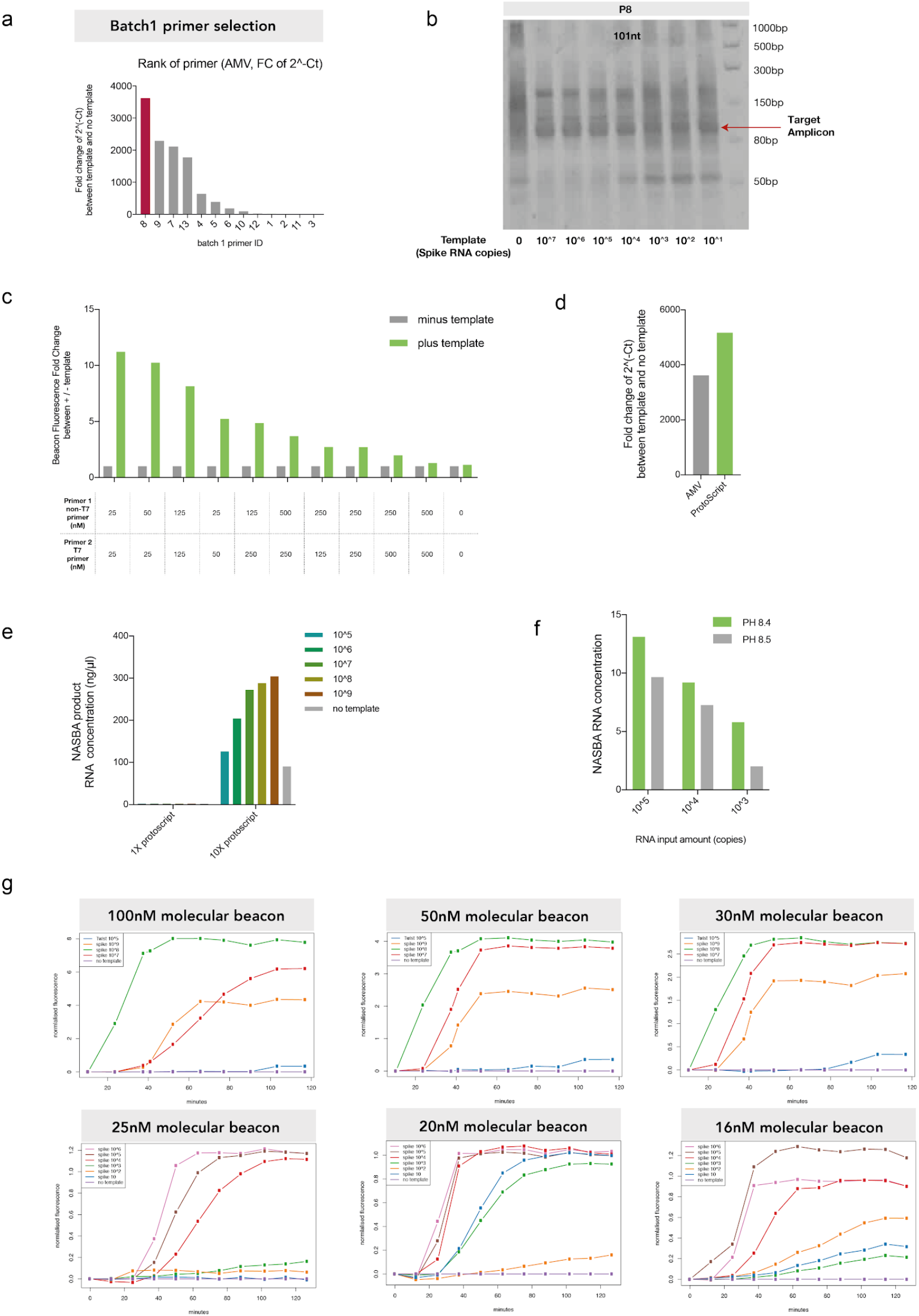
INSIGHT reagent and primer optimisation. (a) Batch 1 primer efficiency test with qPCR as readout. The NASBA end product was used as the template for qPCR with a pair of inner primers. The fold change between the template-containing sample and a sample without template was calculated and ranked. The most efficient primer pair (P8) was used in this work. (b) RNA urea gel for NASBA end product with primer pair P8 and different input template amounts ranging from 10^1^ to 10^7^ copies of viral RNA templates. The amplicon-sized band (101 nt) is indicated with an arrow. (c) High enzyme concentrations improve NASBA efficiency. By adding ten times the amount of ProtoScript II reverse transcriptase, ten times more T7 polymerase and three times increased RNase H, as reported in [21], a significantly higher NASBA end product concentration is detected by the Qubit RNA HS kit at all input template amounts. (d) Primer concentration optimisation. Different amounts of primer concentration were used to test the NASBA reaction efficiency. Previous studies typically used 0.2 - 0.25 μM as a final primer concentration. However, we found ten times lower concentrations resulted in increased NASBA efficiency. The NASBA reaction end-point products were quantified using a molecular beacon. The fold change between template-containing and no-template samples was calculated and used to select the optimal primer concentration. (e) Comparison between two types of reverse transcriptase: AMV and ProtoScript II. The NASBA end product was used as the template for qPCR with a pair of inner primers. The fold change between the template-containing sample and no-template samples was calculated for both enzymes. A higher fold change is observed with ProtoScript II. (f) The NASBA end product RNA concentration at different buffer pH, measured with a Qubit RNA HS assay. A more efficient NASBA reaction was observed with a buffer pH of 8.4 for various amounts of input viral RNA templates. (g) Real-time molecular beacon fluorescence readings for NASBA reaction with different molecular beacon concentrations (16 - 100 nM). The best detection sensitivity was observed with a beacon concentration of 20 nM. The reactions here were carried out with our in-house NASBA mixture.

**Figure S2.**
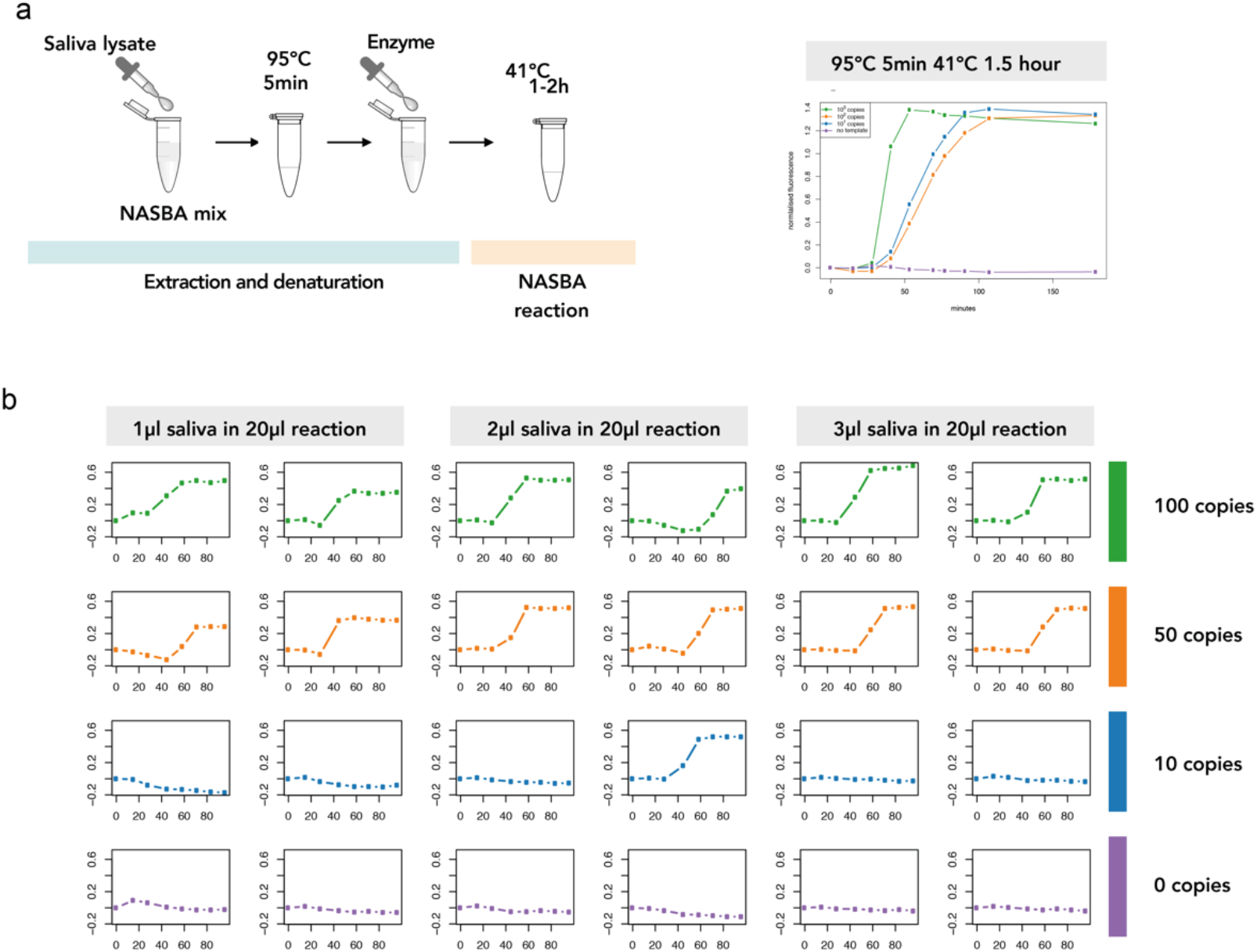
INSIGHT saliva input and optimisation to minimise handling. (a) Left: Protocol of viral lysis in saliva followed by 5 min 95 °C heating before an isothermal NASBA reaction. Right: Real-time molecular beacon fluorescence reading using crude saliva lysate as input, and commercial NASBA mixture under procedures described above, for various amounts of viral RNA input. Barcoded prime pair P8 was used in the reactions. (b) INSIGHT Stage 1 real-time fluorescence readings with various amounts of viral RNA copies (0, 10, 50, 100 copies) and saliva lysate volumes (1 μl, 2 μl and 3 μl) per 20 μl reaction. Reliable detection is achieved for 50 copies of viral RNA input for all saliva input amounts.

**Figure S3.**
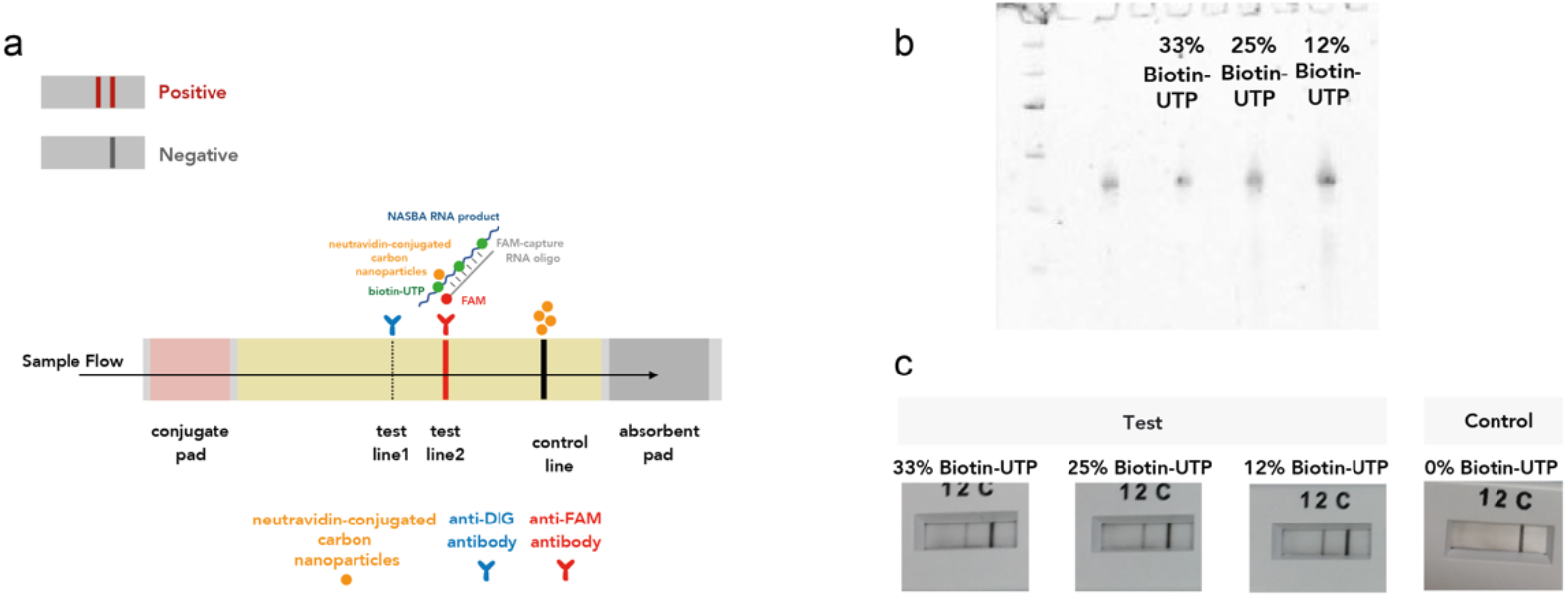
INSIGHT dipstick-based lateral flow assay using biotin-UTP and FAM-labelled oligo. (a) Illustration of the nucleic acid lateral flow assay. Biotinylated UTP can be incorporated into the NASBA RNA products during the reaction. Subsequently, a FAM-labelled RNA capture oligo hybridises to the single-stranded RNA NASBA product. The dual tagging of FAM and biotin results in aggregation of NA-CNP at the test line, resulting in a visible signal in the assay. (b) RNA urea gel for NASBA end product with various proportions (12%, 25%, 33%) of biotin-UTP within the UTP in the reaction. Input template quantity was 10^3^ copies of viral RNA for all three reactions. (c)Lateral flow assay readout of purified RNA from NASBA reactions. Different proportions (12%, 25%, 33%) of biotin-UTP were used. Input template quantity was 10^3^ copies of viral RNA in all three reactions. Here, the NASBA end product needs to be column purified in order to be visualised on the dipstick.

## Supplementary Note: Statistical modelling of barcode hopping

The barcodes here refer to the left and right barcodes used in the NASBA reaction. To illustrate the modelling process, we will use the 48 NASBA reactions carried out in Figure 4b. The left barcodes used in the reactions are denoted as *i*_1_, …, *i*_48_, and the right barcodes as *j*_1_, …, *j*_48_. The used barcode pairs are *i*_1_*j*_1_, *i*_2_*j*_2_, …, *i*_48_*j*_48_, and they are Hamming-distance 3-separated. The DNA concentration of barcode *i*_*n*_*j*_*m*_is denoted as 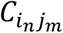, and the read count of barcode *i*_*n*_*j*_*m*_is denoted as 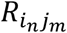.

A barcode pair *i*_*n*_*j*_*m*_ (*n* ≠ *m*) not directly used in the reactions may still have positive read counts. We hypothesise that its reads are mainly a result of 1) barcode hopping and 2) substitution error from any used barcodes that are only Hamming-distance 1 from *i*_*n*_*j*_*m*_.

1. Barcode hopping. We propose that there are 2 mechanisms at play here.

a. Imperfect matching from free floating NASBA primers: residual NASBA (barcoded) primers might bind to imperfectly matched template despite barcode mismatch to start polymerisation in PCR after pooling. This can happen between either template containing barcode pair *i*_*n*_*j*_*n*_ with barcoded primer *j*_*m*_; or between template barcode pair *i*_*m*_*j*_*m*_ with barcoded primer *i*_*n*_.

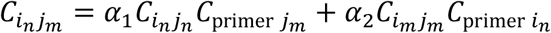 Assuming that concentrations of all free barcoded primers are equal as they exist in excess in NASBA reactions, the above equation can be simplified to:

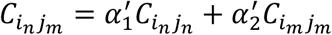
b. Template switching: during PCR, DNA polymerase can jump from one template to another in a region of complementarity, leading to a hybrid sequence [31]. DNA containing *i*_*n*_*j*_*m*_generated in this way should be proportional to the DNA concentration of the 2 original barcoded templates 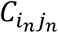 and 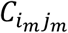.

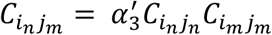
2. Substitution error from any used barcodes that are only Hamming-distance 1 from *i*_*n*_*j*_*m*_:

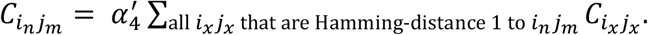 Combining all possible mechanisms:

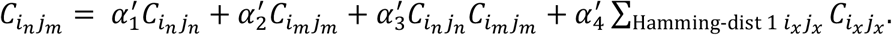 There is an approximately linear relationship between DNA concentration and read count in sequencing, we model 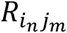 by a negative binomial distribution with mean given by

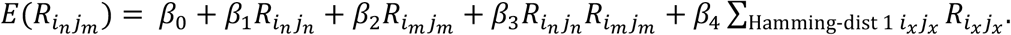 Model is fitted using read counts of all unused barcode pairs *i*_*n*_*j*_*m*_ (*n* ≠ *m*), and the plot below shows that the model can adequately account for almost all such hopped barcode reads.

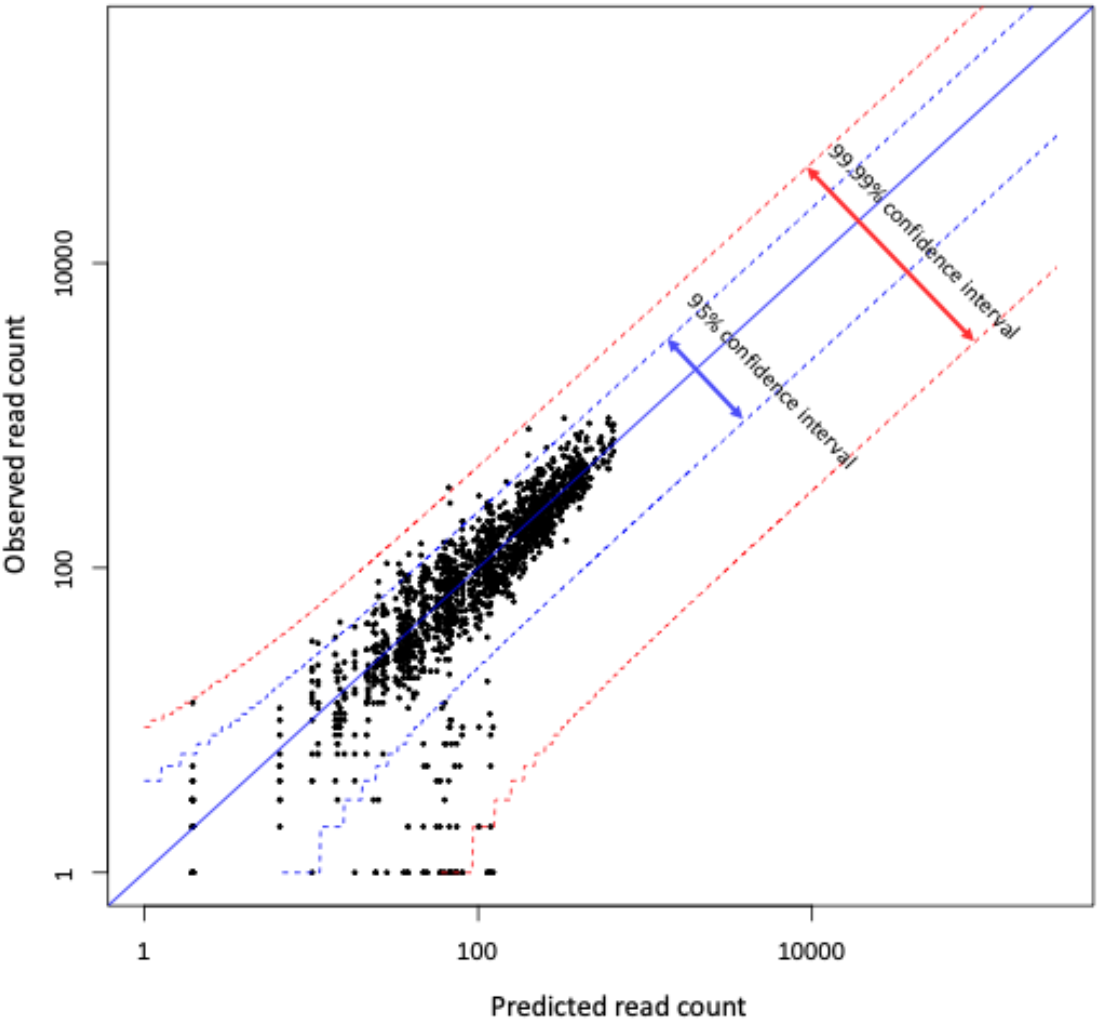

In practice, such a model can be used to distinguish between genuine positive samples from negative samples with reads generated from barcode hopping. By comparing the observed reads with the predicted barcode hopping reads, one can generate Z scores for all mapped barcode pairs. A large Z score means that the read count cannot be solely explained by barcode hopping and substitution error, and is therefore likely to be a result from amplification of a positive sample. We have plotted Z scores for all observed barcode pairs in Figure 4d. In this case, a Z score above 100 can be a clear cut-off for calling of a ‘positive’ sample. The software code for barcode hopping analysis and Z score calculation is available at: *www.github.com/suochenqu/INSIGHT.*

## Supplementary Table

**Table S1.** Composition of in-house NASBA reaction mixture.

Buffer with DMSO
|  | vol. | stock conc. | conc. in RM |
| --- | --- | --- | --- |
| Tris-HCl pH 8.4* | 120 µl | 1 M | 40 mM |
| MgCl<sub>2</sub> | 39.6 µl | 1 M | 13.2 mM |
| KCl | 112.5 µl | 2 M | 75 mM |
| DTT | 30 µl | 1 M | 10 mM |
| DMSO | 450 µl | 100% | 11% |
| water | 247.9 µl |  |  |
| **total volume** | 1000 µl |  |  |

\*Tris-HCl pH 8.4 is made in-house by titrating Tris-HCl pH 8.0 with NaOH pellet and pH determined by pH meter.

Buffer without DMSO
|  | vol. | stock conc. | conc. in RM |
| --- | --- | --- | --- |
| Tris-HCl pH 8.4 | 120 µl | 1 M | 40 mM |
| MgCl<sub>2</sub> | 39.6 µl | 1 M | 13.2 mM |
| KCl | 112.5 µl | 2 M | 75 mM |
| DTT | 30 µl | 1 M | 10 mM |
| water | 697.9 µl |  |  |
| **total volume** | 1000 µl |  |  |

Nucleotide mix (incl. 10% excess)
|  | vol. | stock conc. | conc. in RM |
| --- | --- | --- | --- |
| dNTP | 0.22 µl each | 100 mM | 1 mM each |
| NTP | 0.88 µl each | 100 mM | 4 mM each |
| **total volume** | 4.4 µl |  |  |

Enzyme mix (incl. 10% excess)
|  | vol. | stock conc. | conc. in RM |
| --- | --- | --- | --- |
| diluted RNase H | 0.17 µl | 500 U/ml | 3.75 U/ml |
| ProtoScript RT | 0.28 µl | 200000 U/ml | 2500 U/ml |
| T7 polymerase | 2.75 µl | 50000 U/ml | 6250 U/ml |
| BSA | 0.13 µl | 20 mg/ml | 0.12 mg/ml |
| buffer without DMSO | 1.78 µl |  |  |
| water | 0.40 µl |  |  |
| **total volume** | 5.5 µl |  |  |

**Diluted RNase H**
|  | **vol.** | **stock conc.** |
| --- | --- | --- |
| RNase H | 5 µl | 5000 U/ml |
| BSA (0.48mg/ml) | 1.2 µl | 20 mg/ml |
| buffer without DMSO | 16.67 µl |  |
| water | 27.13 µl |  |
| **total volume** | 50 µl |  |

## Footnotes

2 Molecular beacons were synthesised in the ATDBio laboratory using a K&A H-8 SE DNA Synthesiser, and purified by reversed-phase HPLC.

3 Molecular beacon is reconstituted with annealing buffer (10 mM Tris pH 8 with 10 μM MgCl_2_) to the final concentration of 10 μM. Beacon is then annealed by incubation at 85 °C for 5 min, then gradual cooling to 4 °C by 0.1 °C/s before the NASBA reaction.

## Notes

### Summary of Updates

Methods and Results sections updated details on dipstick readout and sequencing

